# Exercise and angiotensin receptor blockade enhance recovery after orthopaedic trauma in mice by decreasing pain and improving muscle regeneration

**DOI:** 10.1101/773598

**Authors:** Vivianne L. Tawfik, Marco Quarta, Patrick Paine, Thomas E. Forman, Jukka Pajarinen, Yoshinori Takemura, Stuart B. Goodman, Thomas A. Rando, J. David Clark

## Abstract

Chronic pain and disability after limb injury are major public health problems. One key obstacle to addressing these adverse outcomes is that we do not know when exercise should be initiated or whether the beneficial effects of exercise can be reproduced using pharmacological tools. In these studies, we developed and used a murine model of orthopaedic trauma combining tibia fracture and pin fixation with tibialis anterior muscle damage. Behavioral measures included mechanical nociceptive thresholds and distances run on exercise wheels. Bone healing was quantified using microCT scanning, and muscle fiber size distribution as well as fibrosis were followed using immunohistochemistry. We found that the model provided robust mechanical allodynia, fibrosis and a shift to smaller average muscle fiber size lasting up to 5 weeks from injury. We also observed that allowing “late” (weeks 1-2) rather than “early” (weeks 0-1) wheel running after injury resulted in greater overall running activity and greater reversal of allodynia. In parallel, the late running paradigm was also associated with lower levels of muscle fibrosis and a return towards normal muscle fiber diameters. Providing the anti-fibrotic angiotensin receptor blocker losartan to mice in drinking water blocked TGFbeta production while reducing both allodynia and muscle fibrosis. Combining losartan and late exercise provided no additional benefit. We conclude that early healing after orthopaedic trauma must be allowed prior to the initiation of exercise to achieve optimal pain, functional and physiological outcomes. Losartan may provide many of the same pain, functional and physiological outcomes by its regulation of TGFbeta signaling and is a viable candidate for translational studies.

**Key points summary:**

- Our tibial fracture orthopaedic injury model in mice recapitulates the major manifestations of complex trauma including nociceptive sensitization, bone fracture, muscle fibrosis and muscle fiber loss.
- Delayed exercise after complex orthopaedic trauma results in decreased muscle fibrosis and improved pain
- Losartan, an angiotensin-receptor blocker with antifibrotic abilities, recapitulates the effect of exercise on post-injury recovery

## INTRODUCTION

Chronic pain affects 100 million people in the United States alone at an estimated cost of $635 billion/year in medical treatment and lost productivity (Institute of Medicine, 2011). Orthopaedic trauma is a common cause of persistent pain and represents one of five priority areas of the U.S. Bone and Joint Initiative for reducing the burden of musculoskeletal disease (Jacobs et al., 2013). Simultaneous injury to bone, muscle and peripheral nerves make multidisciplinary care (medication optimization, physical therapy, interventional treatments) crucial for patients who sustain orthopaedic trauma. In particular, rehabilitation strategies focused on muscle strengthening and endurance are key to recovery.

After injury, skeletal muscle tissue has a remarkable capacity to regenerate, enabled by resident adult muscle stem cells (MuSCs). However, in pathological conditions, aging or severe traumatic injuries, fibrosis takes over, impeding proper rehabilitation (Baoge et al., 2012; Dueweke, Awan, & Mendias, 2017; Mahdy, 2019). We have previously shown that bioconstructs suffused with MuSCs and other muscle resident cells are effective at treating volumetric muscle loss injuries, an effect that is enhanced when bioconstruct transplantation is followed by exercise (Quarta et al., 2017). Exercise is known to improve analgesia and increase pain thresholds in diverse pain conditions including low back pain, osteoarthritis and myofascial pain (Bement & Sluka, 2016), likely through enhancing central nervous system inhibition and reducing excitation (Fingleton, Smart, & Doody, 2017). In the periphery, exercise increases release of the anti-inflammatory cytokine IL-10 (Leung, Gregory, Allen, & Sluka, 2016), decreases allodynia and paw skin pro-inflammatory cytokines (Shi et al., 2018) and decreases muscle fibrosis by encouraging extracellular matrix remodeling to promote myofiber growth and repair (Garg & Boppart, 2016). However, exercise can also enhance pain (Lima, Abner, & Sluka, 2017) and the exact timing and mechanism of exercise-induced effects in orthotrauma remains unknown.

Angiotensin reception blockers (ARBs), a commonly used and well tolerated class of anti-hypertensive drugs, have been shown to decrease muscle fibrosis and improve muscle strength in several animal models of muscle injury, and mouse and human muscle diseases (Burks et al., 2011; Cohn et al., 2007; Kobayashi et al., 2013). The mechanism is believed to be through the inhibition of TGFbeta1 (Baoge et al., 2012). An interesting possibility is that pharmacologically targeting fibrosis may improve pain outcomes as well. For example, the production of TGFbeta1 has been linked to modulation of inflammatory, visceral, migraine and neuropathic pain, although this modulation may be positive or negative depending on the specific model (Echeverry et al., 2009; Ishizaki et al., 2005; Zhu et al., 2012). With respect to peripherally generated TGFbeta, TGFbeta receptors are expressed on peripheral sensory nerves (Stark, Carlstedt, & Risling, 2001), and TGFbeta1 may enhance nociception through activation of Cdk5 and subsequently TRPV1 receptors (Utreras et al., 2012).

Here we sought to develop an orthopaedic trauma model that recapitulates the major manifestations of such an injury including nociceptive sensitization, bone fracture, muscle fibrosis and muscle fiber loss. Using this model, we tested the hypothesis that exercise enhances recovery after injury as determined by bone callus formation, muscle regeneration and improved mechanical nociceptive thresholds. We further determined whether there was optimal timing to introduce exercise and evaluated the potential beneficial effect of a commonly used ARB drug, losartan, based on its antifibrotic capabilities (Bedair, Karthikeyan, Quintero, Li, & Huard, 2008).

## METHODS

### Ethical Approval

All procedures were approved by the Institutional Animal Care and Use Committee of the Veterans Affairs Palo Alto Healthcare System (Approval #CLA1582) in accordance with American Veterinary Medical Association guidelines and the International Association for the Study of Pain. Male C57BL/6J mice (23-25gr, 11-16 weeks, Jackson Labs) were housed 2-5 per cage before surgery and individually after fracture procedures. Animals were maintained on a 12-hour light/dark cycle in a temperature-controlled environment with *ad libitum* access to food and water.

### Orthopaedic trauma model

Mice were anesthetized with inhalational isoflurane 2-4% and once depth of anesthesia was confirmed with loss of toe pinch reflex, underwent an open right distal tibia fracture followed by intramedullary nail alignment. A skin incision was made from the distal tibia to the proximal tibia to the level of the inferior knee joint on the medial surface. The periosteum was stripped and dried with a cotton swab. A micro drill was then used to make an osteotomy at the proximal end of the tibia at the level of the tibial tuberosity. A 27 G needle was inserted through the osteotomy down the medullary axis of the bone to establish a channel, and subsequently removed. Next, a bone saw was used to score the tibia at the junction of the middle and distal thirds from the lateral aspect causing trauma to the tibialis anterior muscle. The bone fracture was completed using counter pressure. To align the fracture, the 27 G needle was re-inserted into the intramedullary space, through the proximal tibia, and advanced across the fracture site to the distal segment of bone. The needle was then cut flush with the tibial cortex. After hemostasis was confirmed, the wound was closed using a running 5-0 silk suture. Behavioral and other assays began seven days after surgery. After surgery mice received buprenorphine by subcutaneous injection twice daily for 2 days per protocol and were closely monitored throughout the post-anesthesia period and for the length of the study by research personnel.

### Voluntary running

Each subject was singly housed in a cage with a computer controlled running wheel (AWM software, version 6.9.2057.18763; Lafayette Instrument, USA) that was either locked or freely moving. After surgery or behavioral testing, mice assigned to the “early exercise” group were immediately returned to their cages and allowed to run freely for the 7 days before the wheel was locked. Mice in the “delayed exercise” group had their wheels unlocked for 7 days, starting at 7 days post-surgery. “No exercise” controls were singly housed in cages with locked wheels. Each cage wheel was attached to a counter motor and associated data acquisition software that recorded the number of rotations that occurred during each 15 min interval for the length of the study.

### Behavioral testing

To ensure rigor in our findings, all testing was performed by one female experimenter (V.L.T.) who performed all *in vivo* behavioral testing in a blinded fashion to avoid the effect of experimenter sex on outcomes (Sorge et al., 2014). All testing was conducted between 7:00 am - 1:00 pm in an isolated, temperature- and light-controlled room. Mice were acclimated for 30 - 60 min in the testing environment within custom clear plastic cylinders (4” D) on a raised metal mesh platform (24” H). Mice were randomized by simple selection from their home cage prior to testing, and placement in a cylinder, after testing mouse numbers were recorded on the data sheet.

### Mechanical nociception assays

To evaluate mechanical reflexive hypersensitivity^58^, we used a logarithmically increasing set of 8 von Frey filaments (Stoelting), ranging in gram force from 0.007 to 6.0 g. These were applied perpendicular to the plantar hindpaw with sufficient force to cause a slight bending of the filament. A positive response was characterized as a rapid withdrawal of the paw away from the stimulus filament within 4 s. Using the up-down statistical method^70^, the 50% withdrawal mechanical threshold scores were calculated for each mouse and then averaged across the experimental groups.

### Muscle Harvesting & Histology

After euthanasia with inhaled CO_2_, the Tibialis Anterior (TA) muscles were carefully dissected away from the bone, weighed and placed into a 0.5% PFA solution for fixation overnight. The muscles were then moved to a 20% sucrose solution for 3 h or until muscles reached their saturation point and began to sink. The tissues were then embedded and frozen in optimal cutting temperature (OCT) medium and stored at −80°C until sectioning. Sectioning was performed on a Leica CM3050S cryostat that was set to generate 12 μm sections. Sections were mounted on Fisherbrand Colorfrost slides. These slides were stored at −20°C until immunohistochemistry could be performed. For colorimetric staining with Hematoxylin and Eosin (Sigma) or Gomorri Trichrome (Richard-Allan Scientific) samples were processed according to the manufacturer’s recommended protocols. For immunostaining, a 1 h blocking step with 20% donkey serum/0.3% Triton in PBS was used to prevent nonspecific primary antibody binding for all samples. Primary antibodies were applied and allowed to incubate over night at 4**°**C in 20% donkey serum/0.3% Triton in PBS. After four washes with 0.3% PBST, fluorescently conjugated secondary antibodies were added and incubated at room temperature for 1 h in 0.3% PBST. After three additional rinses each slide was mounted using Fluoview mounting media. The following primary antibodies were used: Rabbit anti-Collagen I (Cedarlane Labs, #CL50151AP, 1:200), Rat anti-Laminin (Millipore, #MAB1903, 1:750). Samples were imaged using standard fluorescent microscopy on a Zeiss Axio Observer.Z1 microscope and either a 10X or 20X objective. Volocity imaging software was used to adjust excitation and emission filters and came with pre-programmed AlexaFluor filter settings, which were used whenever possible. All exposure times were optimized during the first round of imaging and then kept constant through all subsequent imaging. Image analysis was conducted using Image J (Schindelin et al., 2012) to calculate the percentage of area composed of collagen by using the color threshold plugin to create a mask of only the area positive for collagen. That area was then divided over the total area of the sample which was found using the free draw tool. All other analyses were performed using Volocity software and manually counting fibers using the free draw tool and also counting the number of nuclei.

### Bone harvesting & histology

After removal of the TA muscle, the tibia was resected from the leg and excess tissue removed and placed immediately in 10% formalin overnight at 4**°**C. Tissue was then washed three times with PBS and placed in 0.5M EDTA at 4**°**C for decalcification. The EDTA was changed every 4-5 days for the next 2 weeks at which time tissue was washed again in PBS three times and transferred to 30% sucrose at 4**°**C for 24 hours. Tissue was then moved to a 50:50 mixture of sucrose and OCT at 4**°**C for 24 h and embedded in OCT. Sectioning was then performed on a Leica cryostat and 10 um sections were directly mounted on Fisherbrand Superfrost Plus Gold slides to minimize detachment. For colorimetric staining with Hematoxylin and Eosin samples were processed according to the manufacturer’s recommended protocols. Sections were visualized on a Zeiss Axio Observer.Z1 microscope with a 10X objective and representative images were taken.

### Computer Tomography protocol

Tibiae were dissected free of soft tissues and the intramedullary nail carefully removed without disrupting the fracture union. The tibiae were then imaged with an eXplore CT120 Micro CT (GE Healthcare, Chicago, IL). The 50 µm scans were performed with 100 kVp, 2 frames per view, 360° rotation, and 2 × 2 binning. Three dimensional reconstructions were made in MicroView (Parallax Innovations, Ontario, Canada), and fracture healing was quantified by determining the volume of bone callus in a consistently-sized region of interest surrounding the central fracture site using Image J (Schindelin et al., 2012).

### Data and statistical analyses

Researchers remained blinded throughout histological, biochemical, and behavioral assessments. Groups were unblinded at the end of each experiment before statistical analysis. Data are expressed as the mean + SD. All data were analyzed using the GraphPad Prism software (version 8.0.0 for Windows, GraphPad Software, San Diego, California USA, www.graphpad.com) with Student’s t tests, or one-way or two-way ANOVA, with a Tukey post-hoc test, as indicated in the main text or figure captions, as appropriate. The number of mice per group is indicated in figure legends and individual data points are provided where appropriate for full data transparency. No mice were excluded from the analyses.

## RESULTS

### Characterization of an orthopaedic trauma model

In order to appropriately develop treatments for complex orthopaedic trauma and surgical repair, we first sought to generate a reproducible mouse model that recapitulates the major manifestations of such an injury including nociceptive sensitization, bone fracture, muscle fibrosis and muscle fiber loss. Based on previous models (Guo et al., 2014), we performed unilateral tibial fracture with intramedullary nail (IMN) stabilization (Fig. 1*A*). As expected, mechanical thresholds decreased significantly after injury and remained significantly lower than baseline out to 5 weeks post-injury (Fig. 1*B*). To further characterize the bone fracture, we performed both micro computed tomography (CT) scans of the fractured limb (Fig. 1*C*) as well as immunohistochemistry on bone sections (Fig. 1*D*) at 4 weeks post-injury. Both methods confirmed irregularities in the cortical bone with callus formation suggesting active repair. Our initial findings suggested that a medial approach to the fracture produced almost no muscle injury (data not shown) while a lateral approach to the fracture generated significant TA fibrosis and fiber loss (Fig. 1*E*-*G*). Fibrosis was clearly increased in the TA at 2 and 4 weeks after injury as evidenced by a visible increase in collagen deposition (Fig. 1*E*). This resulted in a change in overall percent fibrosis from 2.4% in uninjured muscle to 31.5% at 2 weeks and 24.8% at 4 weeks after injury (Fig. 1*F*). Muscle fibers exist in a continuum of sizes with an average diameter of 2000 μM in native murine TA muscles. Regenerating fibers increase in size over time, fully restoring their original size after approximately 4 weeks in physiological conditions (Hardy et al., 2016). However, in our model, we found that regenerating fibers failed to fully increase to a size comparable with uninjured muscles (Fig. 1*G*). Conversely, injured muscles exhibited extensive collagen deposition, suggesting the development of fibrotic tissues that did not resolve even 4 weeks after injury (Fig. 1*G*). These results suggest that our model faithfully recapitulates the key multisystem characteristics of severe orthopaedic trauma and can be used to further explore targeted treatments.

**Figure 1.**
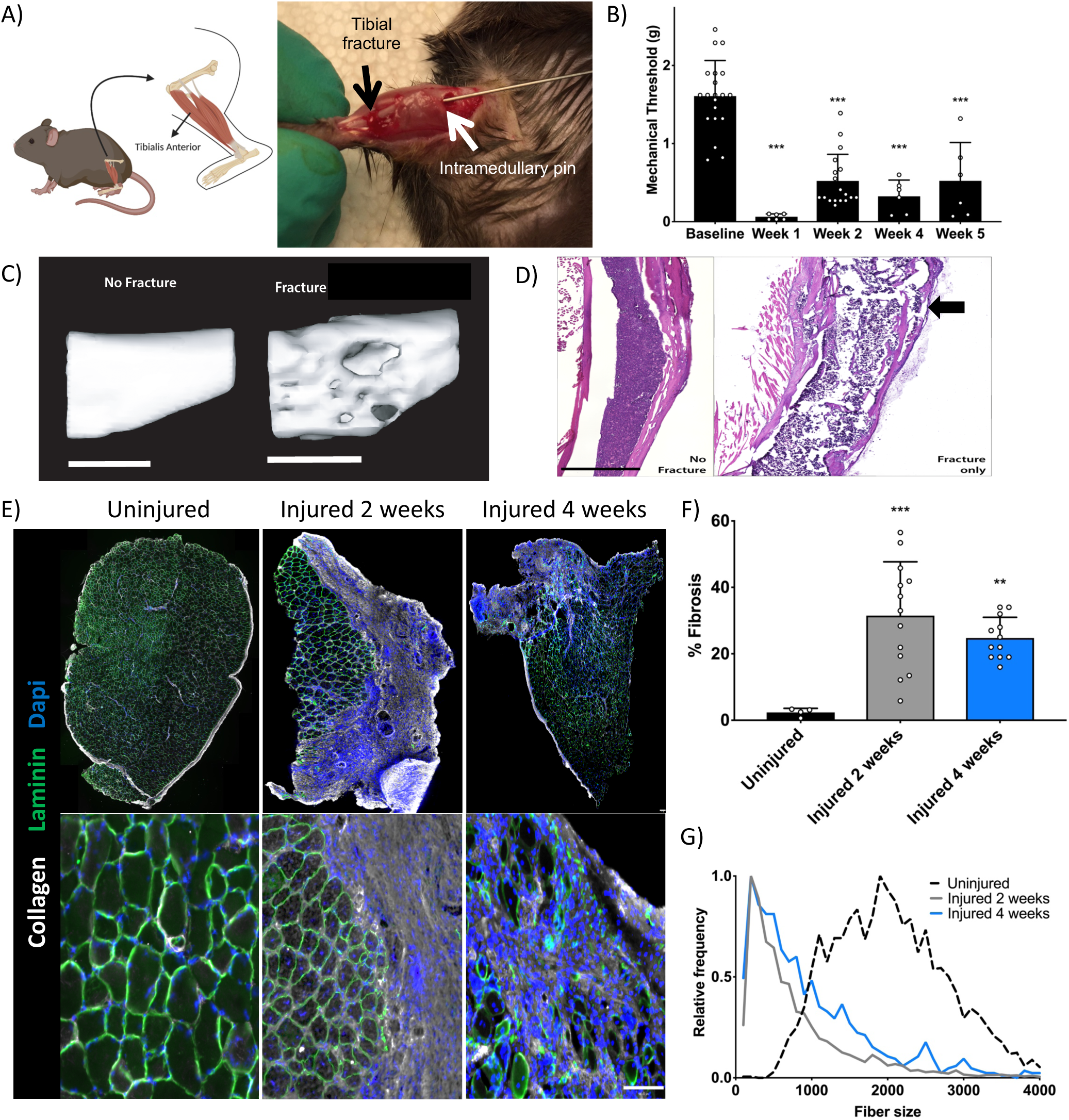
Orthotrauma model recapitulates multisystem injury including nociceptive sensitization, bone fracture, muscle fibrosis and muscle fiber loss. A) Schematic (left, created with BioRender.com) and intra-operative photograph (right) demonstrating location of tibial bone fracture (black arrow), osteotomy and intramedullary nail (white arrow) and tibialis anterior (TA) muscle. B) Mechanical threshold decreases immediately after orthotrauma and remains significantly lower through 5 weeks post-injury (n = 6-20 mice per group, ***p<0.001 vs. uninjured control by one-way ANOVA). C) Micro Computer Tomography (microCT) scan of the right tibia without fracture (left) or 4 weeks after fracture (right) demonstrating clear callus formation. Scale bars 2mm. D) Hematoxylin and Eosin staining of bone sections highlights the cortical disruption that remains at 4 weeks post-injury (black arrow). E) Immunohistochemical staining of TA muscle sections showing increased collagen (white) and altered muscle fiber pattern (laminin, green) as well as loss of regular, central nuclei (DAPI, blue). Scale bars 100 μm. F) Orthotrauma results in extensive fibrosis of the TA muscle (n = 4-13 mice per group, **p<0.01, ***p<0.001 vs. uninjured control by one-way ANOVA). G) Analysis of fiber size frequency shows a clear left shift towards smaller fiber size after fracture that is most pronounced at 2 weeks post-injury. Individual data points are shown in addition to mean + SD for B) and F).

### Delayed exercise improves mechanical allodynia and muscle regeneration

We next investigated whether 7 days of free access to a running wheel either immediately after injury (“early exercise”) or with a one-week delay (“late exercise”), altered behavioral or tissue findings in our orthotrauma model (Fig. 2*A*). Perhaps not surprisingly, mice in the late exercise group that were allowed to recover for one week prior to running wheel access, exercised more than those that had immediate access to a running wheel, as evidenced by a significantly higher total number of wheel rotations (Fig. 2*B*, \*\*\**p*<0.001 early vs late exercise). As has been previously shown (Shi et al., 2018), exercise clearly decreased mechanical sensitivity (Fig. 2*C*). However, this effect was more pronounced and long-lasting in the late exercise group with a return to baseline mechanical threshold at week 4 post-injury (Fig. 2*C*).

**Figure 2.**
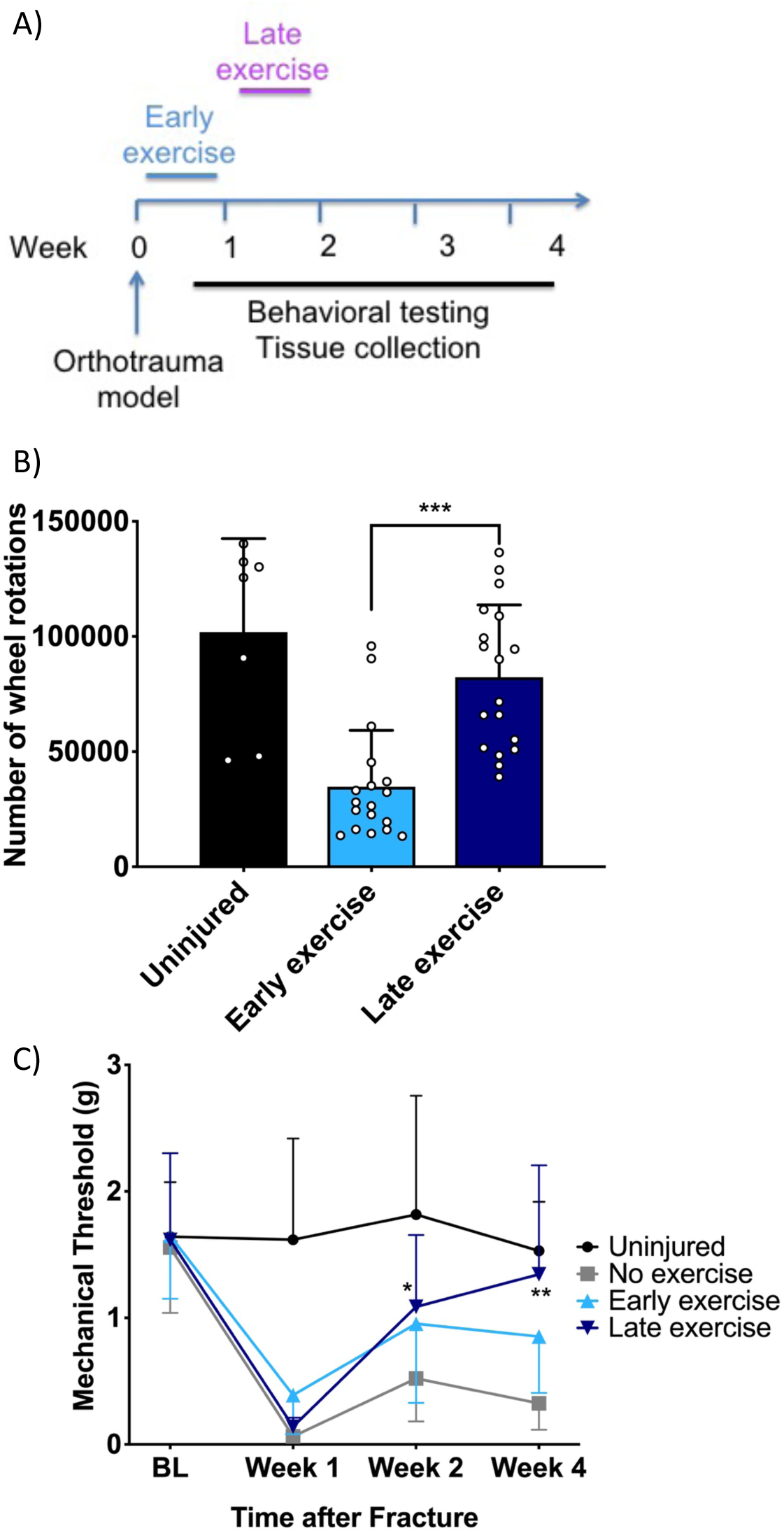
Delayed exercise results in more intensive exercise and improved allodynia trajectory. A) Schematic depicting timeline of exercise with respect to orthotrauma model and measured outcomes. B) The number of wheel rotations mice completed during their 1-week *ad libitum* access to the running wheel (n = 7-18 mice per group, ***p<0.001 vs. early exercise by one-way ANOVA). Individual data points are shown in addition to mean + SD. No data is shown for the “no exercise” fracture group as they did not have access to a functional running wheel. C) The trajectory of mechanical allodynia after fracture is improved by late exercise at week 2 and 4 after injury (n = 6-18 mice per group, *p<0.05, **p<0.01 vs. no exercise by one-way ANOVA).

We next evaluated bone healing using CT scans of the injured lower extremity and found that late exercise reduced the callus size at two weeks and early exercise reduced the callus size at 4 weeks (*SI Appendix*, Fig. S1). We then analyzed the TA muscle for fibrosis and fiber size changes related to exercise. We found that late exercise resulted in a significant decrease in collagen accumulation, and as a result decreased fibrosis, compared to the “no exercise” condition at both 2 and 4 weeks while the early exercise group actually displayed more fibrosis than the “no exercise” group at 4 weeks (Fig. 3*A* and *B*). Most strikingly, there was a prominent increase in fiber size with late exercise at both 2 and 4 weeks with no effect of early exercise on fiber size at all (Fig. 3*C*). Collectively, these data suggest that there is a timeframe during which exercise is most beneficial for bone, muscle and nerve healing.

**Figure 3.**
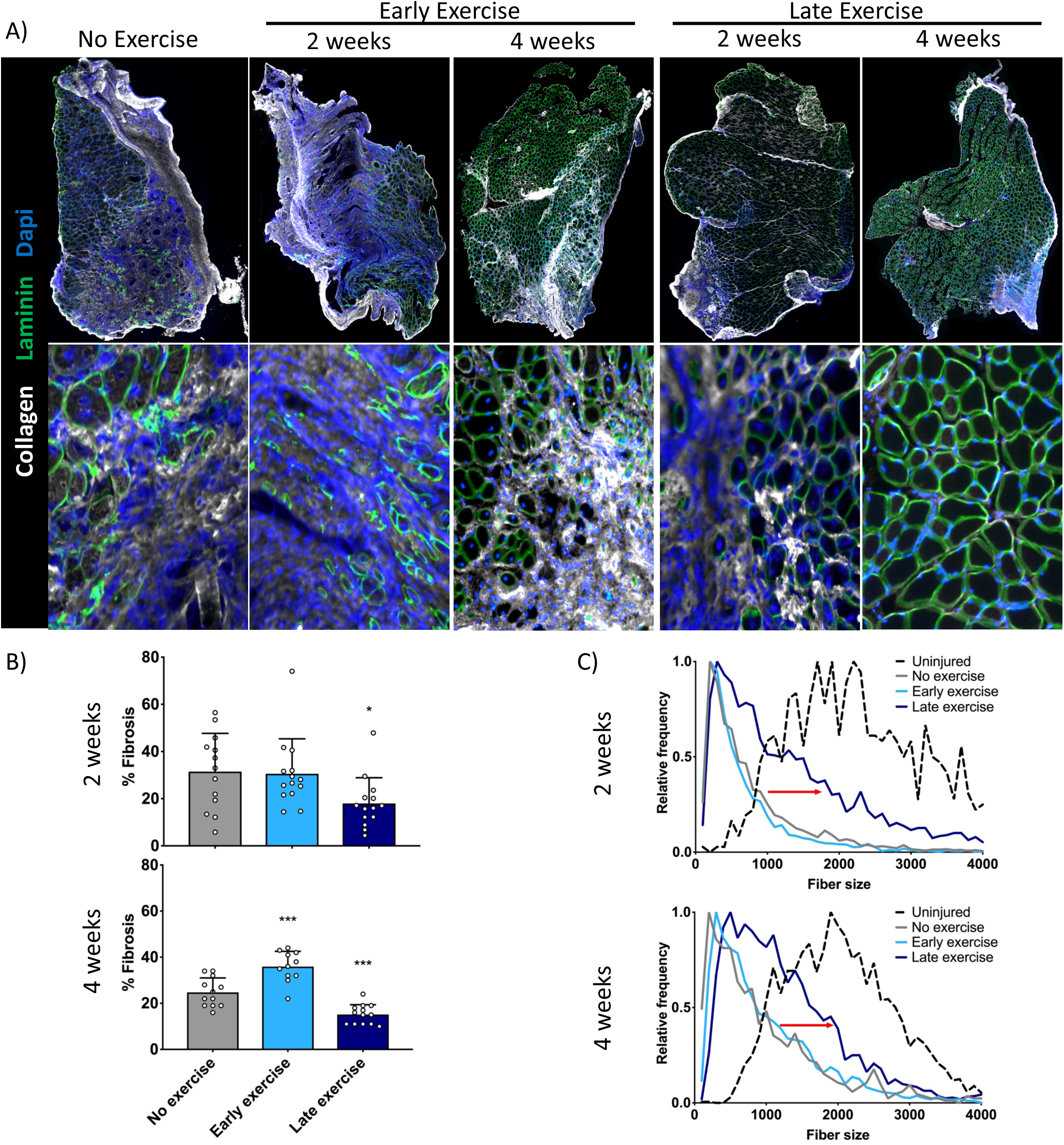
Delayed, but not immediate, exercise improves muscle fiber regeneration and decreases fibrosis. A) Immunohistochemical staining of TA muscle sections shows extensive deposition of collagen (white) and altered muscle fiber pattern (laminin, green) as well as loss of regular, central nuclei (DAPI, blue) after fracture either without exercise or with early exercise. In contrast, TA muscle at either 2- or 4-weeks post-injury had a more organized muscle fiber pattern with low levels of collagen. B) Muscle fibrosis is significantly decreased by late exercise at both 2- and 4-weeks post-injury (n = 12-14 mice per group, *p<0.05, ***p<0.001 vs. no exercise by one-way ANOVA). In contrast, early exercise increases fibrosis at 4 weeks post-injury (n = 12-14 mice per group, ***p<0.001 vs. no exercise by one-way ANOVA). C) Fiber size frequency exhibits a clear right shift towards larger fiber sizes with late exercise at both 2- and 4-weeks post-injury. Individual data points are shown in addition to mean + SD for B).

### Angiotensin receptor blockade recapitulates the effects of late exercise on mechanical allodynia and muscle regeneration

Since fibrosis and muscle fiber regeneration were dramatically improved by late exercise, we sought to determine whether these results could be replicated using a pharmacologic approach. Previous studies have demonstrated an antifibrotic effect of angiotensin II receptor type 1 (AT1R) antagonism in other models of muscle injury (Bedair et al., 2008), and we therefore selected the well tolerated AT1R blocker, losartan, to pursue additional studies. Losartan was administered in drinking water immediately after injury for 4 weeks, either alone or in combination with late exercise (Fig. 4*A*). We first compared the number of wheel rotations performed by mice in the late exercise + losartan group with mice (from Fig. 2*B*) that were treated with late exercise only and found no difference between groups (Fig. 4*B*). We next evaluated the effect of losartan administration on mechanical allodynia, with or without concomitant exercise, and found that treatment resulted in a near complete prevention of allodynia (Fig. 4*C*). Losartan treatment did not have any effect on bony callus volume (*SI Appendix*, Fig. S2). However, consistent with prior studies (Bedair et al., 2008; Kobayashi et al., 2013), losartan alone resulted in decreased muscle fibrosis (Fig. 5*A* and *B*) and a right-shift in fiber size suggestive of improved muscle fiber regeneration (Fig. 5*C*). Finally, we confirmed that this was a process correlated with TGF-β as the levels of this pro-fibrotic cytokine were significantly decreased in TA muscle after treatment with losartan, with or without exercise, but not from exercise alone (Fig. 5*D* and *E*).

**Figure 4.**
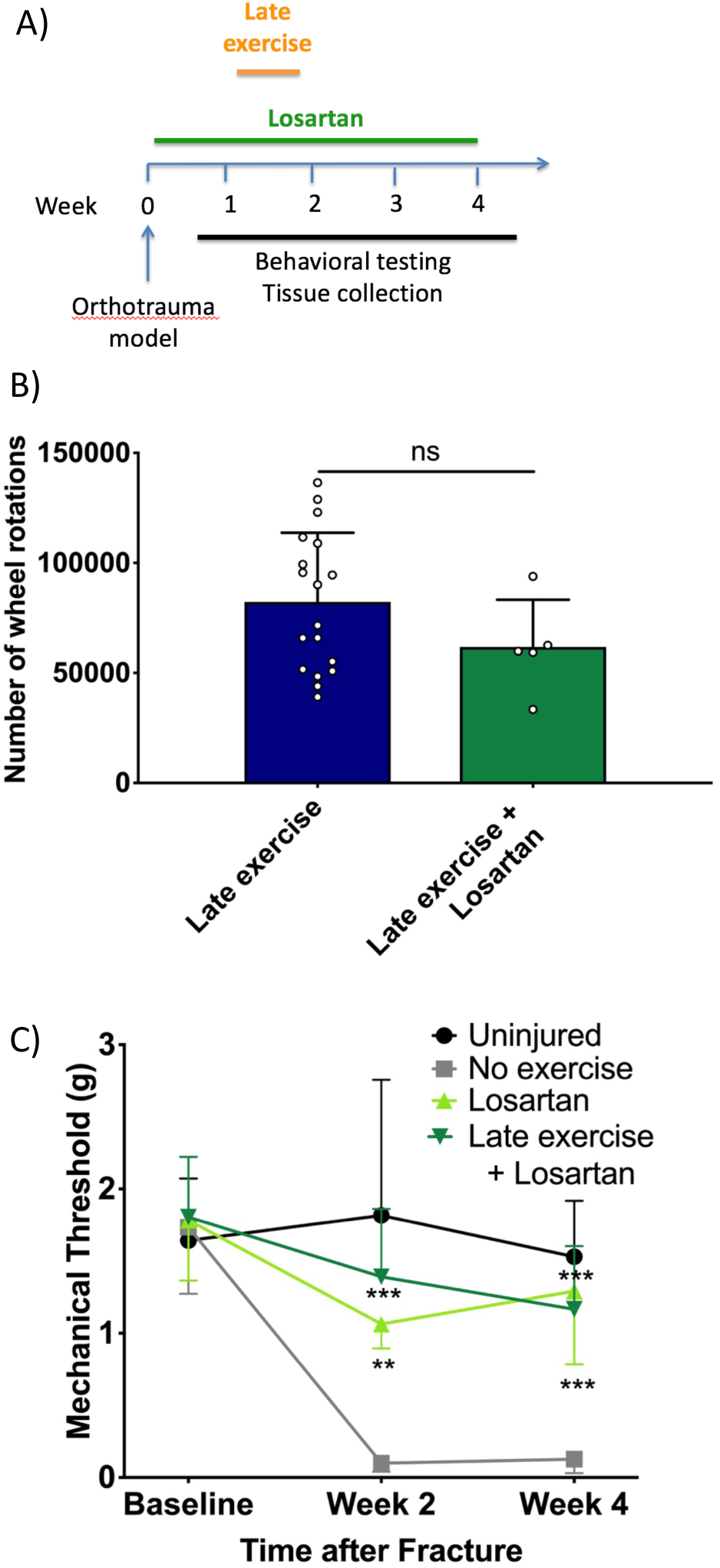
The angiotensin receptor blocker, losartan, improves allodynia after orthotrauma. A) Schematic depicting timeline of exercise and losartan treatment with respect to orthotrauma model and measured outcomes. B) The number of wheel rotations mice completed during their 1-week *ad libitum* access to the running wheel was not significantly different between the late exercise and late exercise + losartan groups (n = 5-18 mice per group, p>0.05 by one-way ANOVA). Individual data points are shown in addition to mean + SD. No data is shown for the losartan only group as they did not have access to a functional running wheel. C) The trajectory of mechanical allodynia after fracture is improved by losartan alone or late exercise + losartan at 2- and 4-weeks after injury (n = 5-12 mice per group, **p<0.01, ***p<0.001 vs. no exercise by two-way ANOVA), however, no additive effect of losartan on exercise was appreciated (losartan vs. late exercise + losartan, p>0.05 by two-way ANOVA).

**Figure 5.**
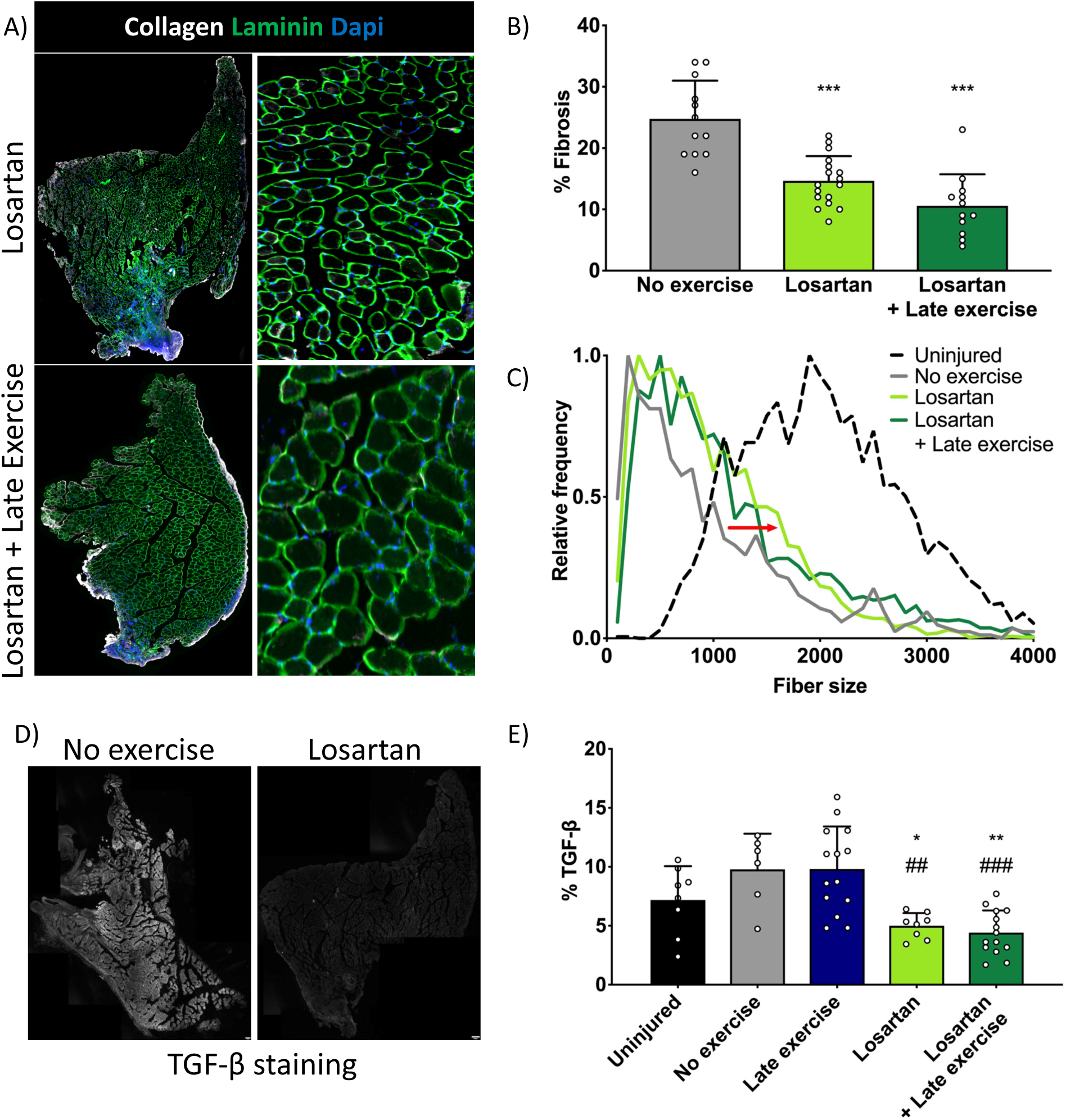
Losartan improves muscle regeneration, decreases fibrosis and suppresses TGF-β expression. A) Immunohistochemical staining of TA muscle sections shows improved collagen deposition (white) and more regular muscle fiber pattern (laminin, green) and central nuclei (DAPI, blue) after losartan treatment either with or without exercise at 4 weeks post-injury. B) Muscle fibrosis is significantly decreased at 4 weeks by losartan treatment with or without late exercise (n = 12-17 mice per group, ***p<0.001 vs. no exercise by one-way ANOVA). C) Fiber size frequency exhibits a clear rightward shift towards larger fiber sizes with losartan with or without late exercise at 4 weeks post-injury. D) Immunohistochemical staining of TA muscle sections with TGF-β antibody demonstrates post-fracture increase in staining that is suppressed by losartan treatment at 4 weeks post-injury. E) %TGF-β staining is significantly decreased by losartan treatment either with or without exercise but not by late exercise alone (n = 6-14 mice per group, *p<0.05, **p<0.01 vs. no exercise; ^##^p<0.01, ^###^p<0.001 vs. late exercise by one-way ANOVA). Individual data points are shown in addition to mean + SD for B) and E).

## DISCUSSION

Orthopaedic injuries involving bone and muscle trauma are common in many settings including falls, motor vehicle accidents and military combat. In this study we demonstrate that 1) muscle fibrosis contributes to poor recovery after orthopaedic trauma, 2) delayed exercise improves muscle regeneration and enhances analgesia, and 3) these effects can be recapitulated using the commonly prescribed angiotensin receptor blocker, losartan. These findings are of both potential scientific and clinical value with particular significance in light of the nation’s ongoing opioid crisis. Approaches to rehabilitation using exercise, non-opioid medications or combinations of approaches may reduce reliance on opioids during the recovery phase after common orthopaedic injuries and surgery. Current evidence suggests that approximately 6% of those patients receiving opioids after acute injuries and surgeries go on to long-term use (Brummett et al., 2017). Our findings both refine our understanding of the importance of timing of rehabilitation for most rapid pain resolution and functional recovery, and also suggest that modulating TGF-beta signaling may be of value to those not able to fully participate in exercise therapy after injuries.

An important finding from our studies is that introducing exercise immediately after orthopaedic trauma is not beneficial and may actually worsen recovery. There are alternative explanations for this observation. One possibility is that the effect of exercise is dose-dependent. Mice given early access to the running wheel actually performed fewer wheel rotations in the same time frame as those given delayed access to the running wheel. Interestingly, however, early exercise mice actually exhibited worse fibrosis compared to control (non-exercised) mice and no beneficial effect was seen on bone callus formation and fracture healing (Figure 3 and Supp Figure 2). This could have occurred because of high strain and repeated microdamage to the fracture site on an injury that was still unstable (Einhorn & Gerstenfeld, 2015). Another explanation for these findings is that immediately after injury, the pro-inflammatory milieu is critical to proper healing (Loi, Cordova, Pajarinen, et al., 2016). In agreement with this theory, we have shown that too early conversion of M1 pro-inflammatory to M2 anti-inflammatory macrophages is detrimental to bone healing as acute inflammation is required for osteogenesis promoted by mesenchymal stem cells and osteoprogenitors (Loi, Cordova, Zhang, et al., 2016). In relation to exercise, in the tibia fracture cast model we found that it at least transiently reduced cytokine expression in injured limbs (Shi et al., 2018) and furthermore exercise enhances muscle M2 macrophages (Leung et al., 2016). It is therefore possible that too early a resolution of the post-traumatic inflammatory state is actually detrimental to muscle regeneration because of alteration in cytokines, such as IL-10, that promote muscle growth and regeneration through immunomodulatory mechanisms (Deng, Wehling-Henricks, Villalta, Wang, & Tidball, 2012; Tidball, 2017; Tidball & Villalta, 2010).

Our findings clearly highlight the potential of ARBs as agents with analgesic potential in conditions where fibrosis is central to the injury and may serve as a trigger for continued pain. Interestingly, in human studies the use of ARBs has been associated with improved muscle strength in patients undergoing chronic hemodialysis (Lin et al., 2019). Such findings have been confirmed in a variety of pre-clinical muscle injury models. For example, in a model of direct gastrocnemius injury/laceration, losartan increased the number of regenerating fibers and decreased the amount of fibrosis in the muscle (Bedair et al., 2008). These findings were extended to a muscle contusion model in which losartan, dosed either immediately at the time of injury or in a delayed manner, improved muscle fibrosis and regeneration (Kobayashi et al., 2013). Furthermore, secondary muscle damage after carbon tetrachloride-induced hepatotoxicity is reduced by losartan via a direct effect on TGF-B1 levels (Hwang et al., 2016). Moreover, losartan has been tested in preclinical models as a strategy to improve recovery from several traumatic injuries, such as volumetric muscle loss (Garg, Corona, & Walters, 2014). In contrast to our findings, however, the latter study did not demonstrate any muscle regeneration with losartan treatment, perhaps because the muscle loss in the VML injury is more extensive than in our current orthopaedic trauma model. Finally, losartan has also been shown to normalize muscle architecture and improve muscle function in the *fibrillin-1*-deficient mouse, a model of the connective tissue disease, Marfan’s syndrome, and in *mdx* mice, a model of Duchenne muscular dystrophy (Cohn et al., 2007). Our own findings in a lower extremity orthopaedic trauma model confirm the ability of losartan to decrease muscle fibrosis and increase regeneration in a TGF-B dependent manner. Most importantly, however, we further demonstrate a potent analgesic effect of losartan in our model, effectively reversing existing allodynia. When losartan was combined with exercise, we did not see a synergistic or even additive effect of the interventions. This suggests that the mechanism of exercise-induced analgesia and muscle regeneration is heavily dependent on AT1R receptor activation and downstream signaling molecules such as TGF-B. Importantly, however, direct inhibition of TGF-B provides early muscle recovery but possible more complex long-term changes such as irregularity of regenerated muscle that may be detrimental (Gumucio, Flood, Phan, Brooks, & Mendias, 2013). The efficacy of TGF-B directed therapy thus warrants further study to more fully understand potential side effects.

The pharmacologic target of ARBs is the renin-angiotensin system, known to regulate blood pressure and electrolyte balance; however, it has also been implicated in the modulation of pain (Bessaguet, Magy, Desmouliere, & Demiot, 2016). For example, intrathecal angiotensin II is sufficient to produce nociceptive behaviors (scratching, biting, licking) which can be reversed by co-administration of intrathecal losartan (Nemoto et al., 2013). Additionally, in several animal models of pain including chronic constriction injury of the sciatic nerve (Pavel, Oroszova, Hricova, & Lukacova, 2013) and chemotherapy-induced neuropathic pain (Kim, Hwang, Kim, Abdi, & Kim, 2019), AT1R blockade decreased allodynia through suppression of primary afferent proinflammatory cytokine expression. These findings suggest that apart from antifibrotic capability of AT1R antagonism, antinociceptive effects of ARBs may arise from direct effects on sensory neurons (Figure 6C).

**Figure 6.**
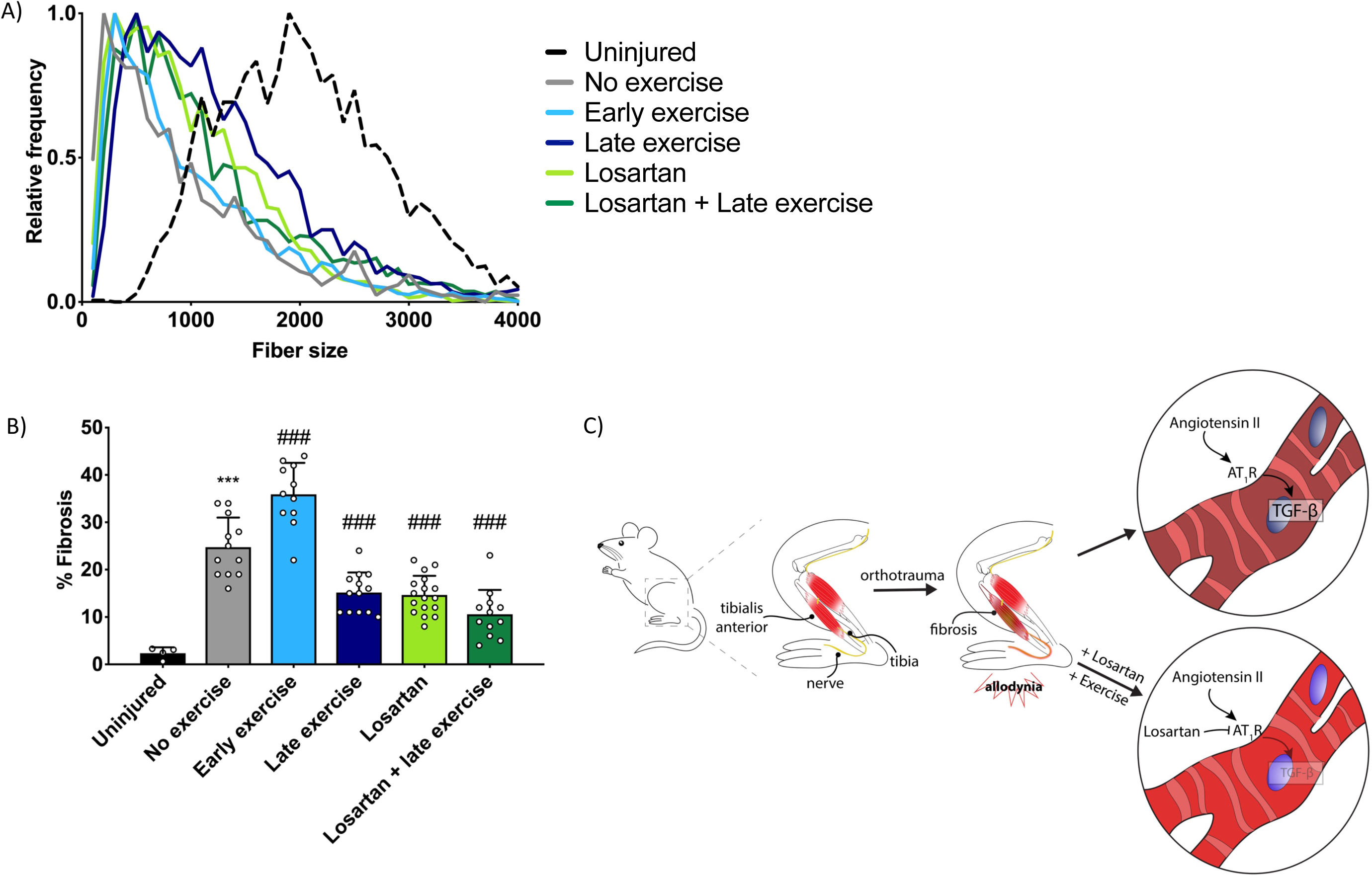
Summary of interventions and proposed mechanism. A) Muscle regeneration as indicated by increased frequency of larger fiber sizes is most apparent with losartan, late exercise or late exercise + losartan. B) Muscle fibrosis is exacerbated by early exercise and improved by losartan, late exercise or late exercise + losartan (n = 4-17 mice per group, ***p<0.001 no exercise vs. uninjured, ^###^p<0.001 no exercise vs. all other groups by one-way ANOVA). Individual data points are shown in addition to mean + SD. C) Schematic outlining the proposed mechanisms of action of losartan and exercise on muscle fibrosis.

At this time the treatment of chronic pain is largely based on general algorithms and medications trials to determine efficacy of a given medication in an individual patient. Moving towards mechanism-based treatment is key to improving the care of patients with chronic pain. A well-tolerated FDA-approved drug such as losartan may prove to have benefit in patients with underlying irrecoverable muscle injuries and degenerative processes driven by fibrosis and chronic inflammation, as a consequence of trauma or aging. Future translational studies evaluating this approach are warranted to determine the treatment implications of our findings.

## ACKNOWLEDGMENTS

This work was supported by the Veterans Affairs RR&D Merit Grant Award I01RX001475 (J.D.C.), the Veterans Affairs RR&D Merit Review I01RX001222 (T.A.R.), the National Institutes of Health Grant R01NS088339 (J.D.C.), the National Institutes of Health (NIH) Grant K08NS094547 (V.L.T.), the Foundation for Anesthesia Education and Research (FAER) Mentored Research Training Grant (V.L.T.) as well as funding from the University of Toyama (Y.T.). The authors wish to thank the technicians in the Veterinary Medical Unit at the Veterans Affairs Hospital Palo Alto Branch for their assistance.

## AUTHOR CONTRIBUTIONS

V.L.T. performed behavioral studies. J.P., T.E.F., P.P., M.Q. and V.L.T. performed CT scans. Y.T., T.E.F., P.P., M.Q. and V.L.T. performed histology. V.L.T, M.Q. and J.D.C. designed studies and wrote the manuscript. All authors contributed to data analysis and editing of the manuscript. S.B.G. designed and supervised the CT scans. T.A.R. and J.D.C. conceived the project, provided critical input on study design and interpretation and supervised all experiments. All authors approved the final version of this manuscript, agree to be accountable for all aspects of this work and agree that all those listed qualify for authorship.

## Supplemental Figure Legends

**Supplemental Figure 1. Size of fracture callus is changed with exercise.** Micro A) Computer Tomography (microCT) scans of the right tibia without fracture (left) or at 2 weeks (top row) or 4 weeks (bottom row) after fracture with and without exercise demonstrating clear callus formation. B) Volumetric analysis of callus formation demonstrated that callus volume was similar in the “no fracture” control and the fracture late exercise group at 2 weeks and the fracture early exercise group at 4 weeks (n = 5-6 per group, p>0.05). Scale bars 2mm.

**Supplemental Figure 2. Histological evaluation of bone microhealing shows no effect of losartan on bony callus area.** A) Sections of fractured tibia were stained with hematoxylin and eosin to evaluate the bony callus at 4 weeks post-fracture. B) No difference was seen between groups with respect to callus area. n = 4-5 mice per group. Scale bars 2mm.

