## Supplemental figures 1&2 for "Exercise and angiotensin receptor blockade enhance recovery after orthopaedic trauma in mice by decreasing pain and improving muscle regeneration" (from another version of this preprint: version 1, 2019-09-18)

### Supplemental 1: Size of fracture callus is decreased with exercise

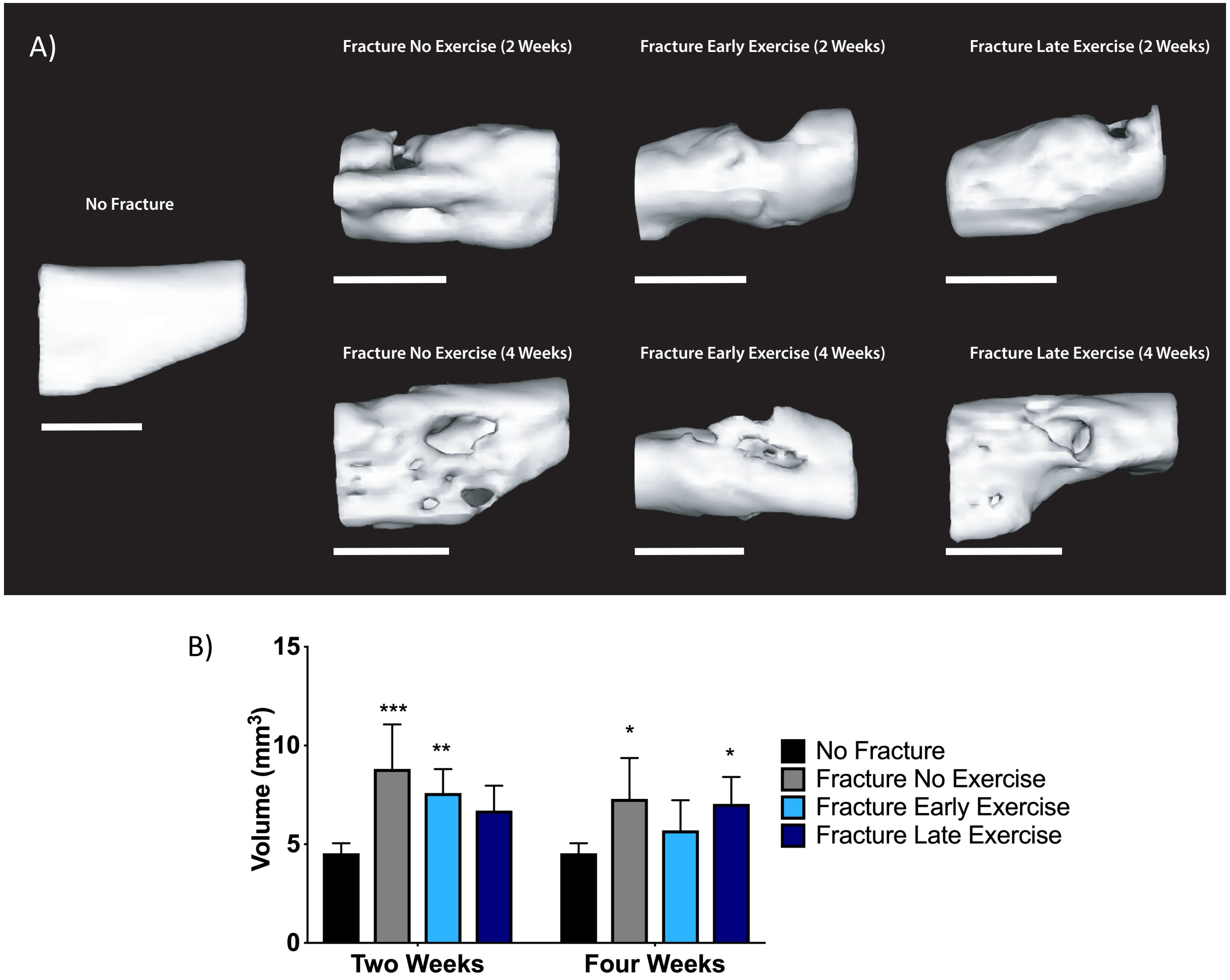

### Supplemental 2: Bone microhealing is unchanged by losartan treatment

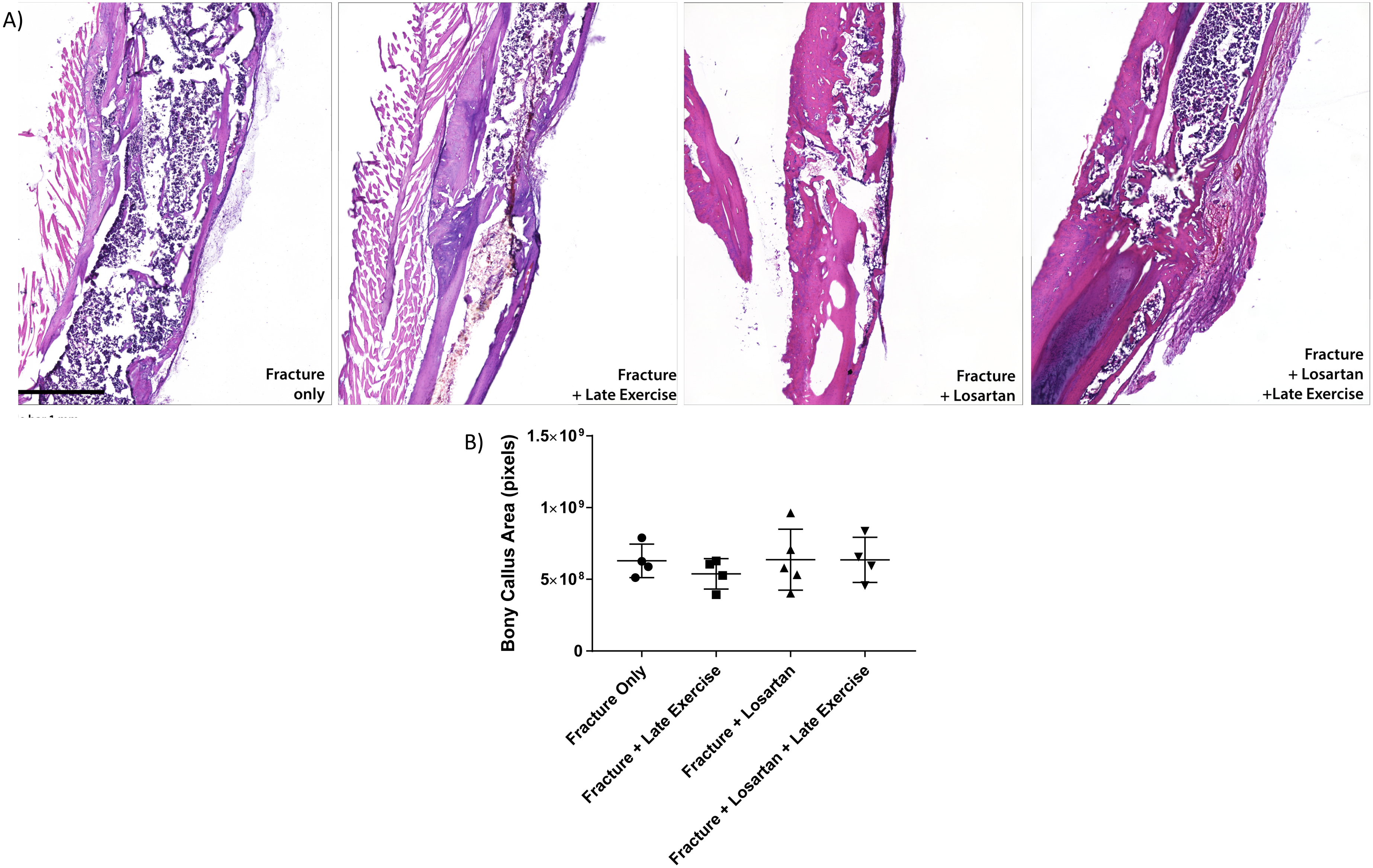
